## Supplemental tables and figures for "Behaviourally modulated hippocampal theta oscillations in the ferret persist during both locomotion and immobility"

| Animal ID | N sessions | Mean session time (minutes ± SD) | Mean trials per session (N ± SD) |
|---|---|---|---|
| R1 | 27 | 69 ± 22 | 65 ± 44 |
| R2 | 15 | 45 ± 17 | 58 ± 39 |
| R3 | 13 | 46 ± 18 | 56 ± 49 |
| F1 | 20<br>(L: 12 ,R: 8) | 35 ± 6 | 50 ± 15 |
| F2 | 23<br>(L: 23) | 33 ± 9 | 45 ± 11 |
| F3 | 32<br>(L: 10,R: 22) | 32 ± 10 | 42 ±19 |

**Supplementary Table 1: Number and length of recording sessions for each animal**

| | Full | Rat | Ferret |
|---|---|---|---|
| Movement (Immobile\|Moving) | 0.017 (0.015, 0.019) t = 19.489 p < 0.001 | 0.190 (0.189, 0.192) t = 270.808 p < 0.001 | 0.017 (0.015, 0.019) t = 19.295 p < 0.001 |
| Species (Ferret\|Rat) | -0.184 (-0.280, -0.088) t = -3.753 p = 0.0002 | | |
| Channel (SO\|SR/SLM) | 0.067 (0.066, 0.068) t = 121.754 p < 0.001 | 0.012 (0.011, 0.014) t = 17.566 p < 0.001 | 0.136 (0.134, 0.138) t = 163.140 p < 0.001 |
| Movement * Species | 0.174 (0.171, 0.176) t = 152.014 p < 0.001 | | |
| Constant | 0.427 (0.359, 0.495) t = 12.325 p < 0.001 | 0.261 (0.188, 0.333) t = 7.043 p < 0.001 | 0.408 (0.312, 0.505) t = 8.272 p < 0.001 |

**Supplementary Table 2: Linear mixed-effects model results predicting peak range across species and movement contingencies (for data in Figure 4)**
Numbers in brackets represent 95% confidence intervals. SO: stratum oriens, SR/SLM: stratum radiatum/stratum lacunosum moleculare.

| | Full | Moving | Immobile |
|---|---|---|---|
| Movement (Immobile\|Moving) | 0.122<br>(0.117, 0.127)<br>t = 47.931<br>p < 0.001 | | |
| Drug Condition (Atropine\|Control) | 0.129<br>(0.117,0.140)<br>t = 21.308<br>p < 0.001 | -0.007<br>(-0.033, 0.020)<br>t = -0.483<br>p = 0.630 | 0.129<br>(0.116, 0.141)<br>t = 20.297<br>p < 0.001 |
| Movement * Drug Condition | -0.139<br>(-0.147, -0.132)<br>t = -37.271<br>p < 0.001 | | |
| Constant | 0.339<br>(0.311, 0.367)<br>t = 23.668<br>p < 0.001 | 0.453<br>(0.417, 0.489)<br>t = 24.691<br>p < 0.001 | 0.340<br>(0.310, 0.370)<br>t = 22.387<br>p < 0.001 |

**Supplementary Table 3: Linear mixed-effects model results predicting peak range in ferrets across drug and movement contingencies (for data in Figure 5)**
Numbers in brackets represent 95% confidence intervals.

| | Full rat | SO | SR/SLM |
|---|---|---|---|
| Trial Epoch (Hold\|Run) | -0.285 (-0.292, -0.277) t = -72.775 p < 0.001 | -0.285 (-0.292, -0.277) t = -74.072 p < 0.001 | -0.211 (-0.218, -0.203) t = -53.369 p < 0.001 |
| Trial Epoch (Hold\|Reward) | -0.311 (-0.319, -0.303) t = -79.535 p < 0.001 | -0.311 (-0.319, -0.304) t = -80.953 p < 0.001 | -0.233 (-0.241, -0.226) t = -59.103 p < 0.001 |
| Channel (SO\|SR/SLM) | -0.057 (-0.065, -0.049) t = -13.694 p < 0.001 | | |
| Hold * Channel | 0.074 (0.063, 0.085) t = 12.717 p < 0.001 | | |
| Reward * Channel | 0.078 (0.066, 0.089) t = 13.372 p < 0.001 | | |
| Constant | 0.553 (0.473, 0.634) t = 13.515 p < 0.001 | 0.541 (0.479, 0.603) t = 17.116 p < 0.001 | 0.554 (0.429, 0.679) t = 8.690 p < 0.001 |

**Supplementary Table 4: Linear mixed-effects model results predicting peak range in rats across trial epochs (for data in Figure 6)**
Numbers in brackets represent 95% confidence intervals. SO: stratum oriens, SR/SLM: stratum radiatum/stratum lacunosum moleculare.

| | Full ferret | SO | SR/SLM |
|---|---|---|---|
| Trial Epoch (Hold\|Run) | -0.035 (-0.042, -0.028) t = -9.787 p < 0.001 | -0.035 (-0.042, -0.029) t = -10.786 p < 0.001 | -0.087 (-0.095, -0.078) t = -20.237 p < 0.001 |
| Trial Epoch (Hold\|Reward) | -0.030 (-0.037, -0.023) t = -8.364 p < 0.001 | -0.030 (-0.037, -0.024) t = -9.218 p < 0.001 | 0.107 (0.098, 0.115) t = 24.823 p < 0.001 |
| Channel (SO\|SLR/SLM) | 0.119 (0.111, 0.127) t = 28.710 p < 0.001 | | |
| Hold * Channel | -0.052 (-0.063, -0.040) t = -9.067 p < 0.001 | | |
| Reward * Channel | 0.137 (0.126, 0.148) t = 24.071 p < 0.001 | | |
| Constant | 0.449 (0.335, 0.562) t = 7.751 p < 0.001 | 0.447 (0.346, 0.549) t = 8.642 p < 0.001 | 0.497 (0.388, 0.607) t = 8.920 p < 0.001 |

**Supplementary Table 5: Linear mixed-effects model results predicting peak range in ferrets across trial epochs (for data in Figure 6)**
Numbers in brackets represent 95% confidence intervals. SO: stratum oriens, SR/SLM: stratum radiatum/stratum lacunosum moleculare.

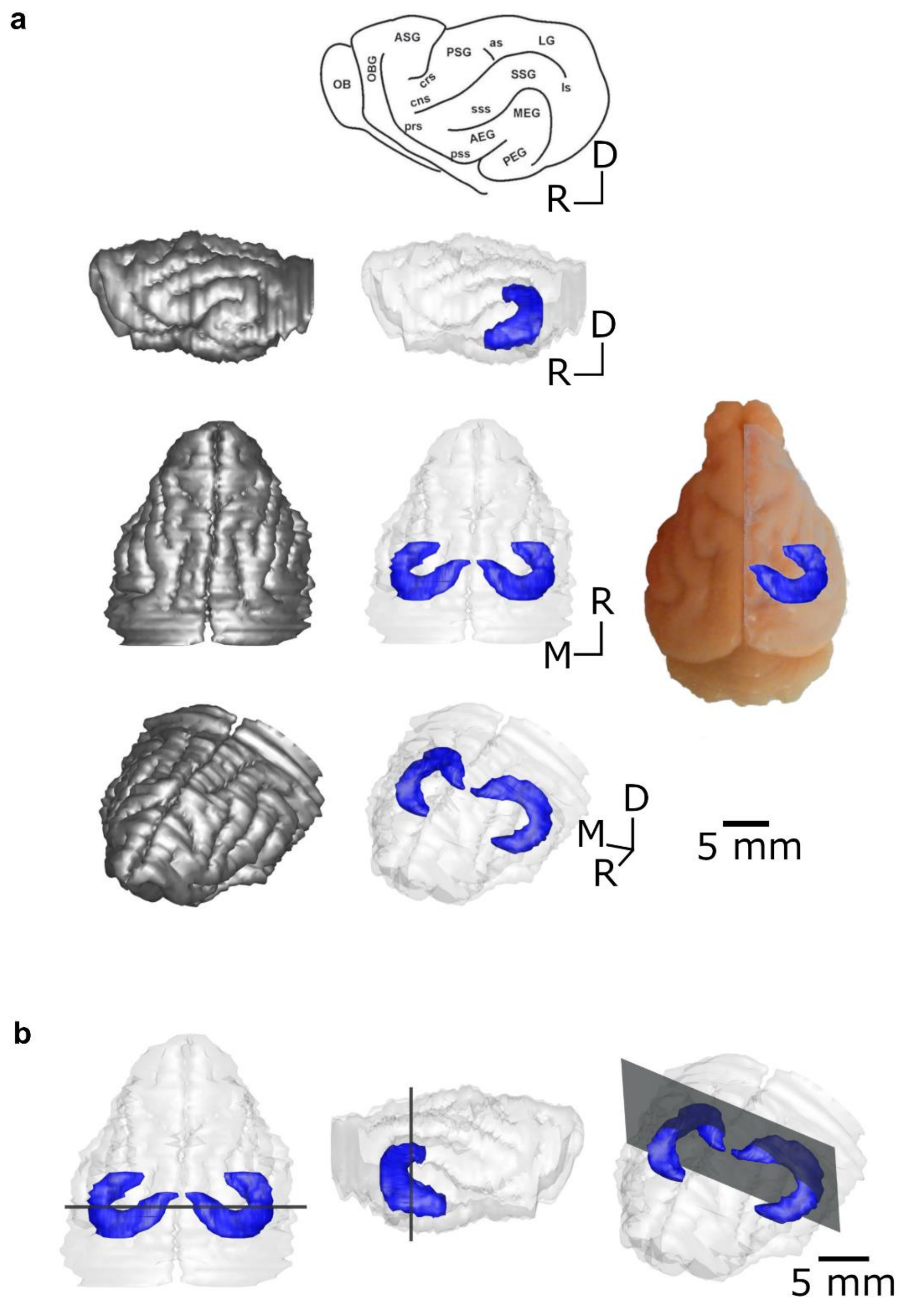


**Extended Data Figure 1: Gross anatomy of the ferret hippocampus**

**a,** Schematic of the ferret brain. 3D reconstruction of the ferret brain (reconstructed from ferret F1). Left: cortical surface; middle: position of the hippocampus within the ferret brain; right: 3D model overlaid on image of the brain used for the reconstruction. **b,** 3D reconstruction of the brain of ferret F1, indicating the location of the histological slice in Figure 1b with grey planes (Left: top view, middle: side view, right: angled view).

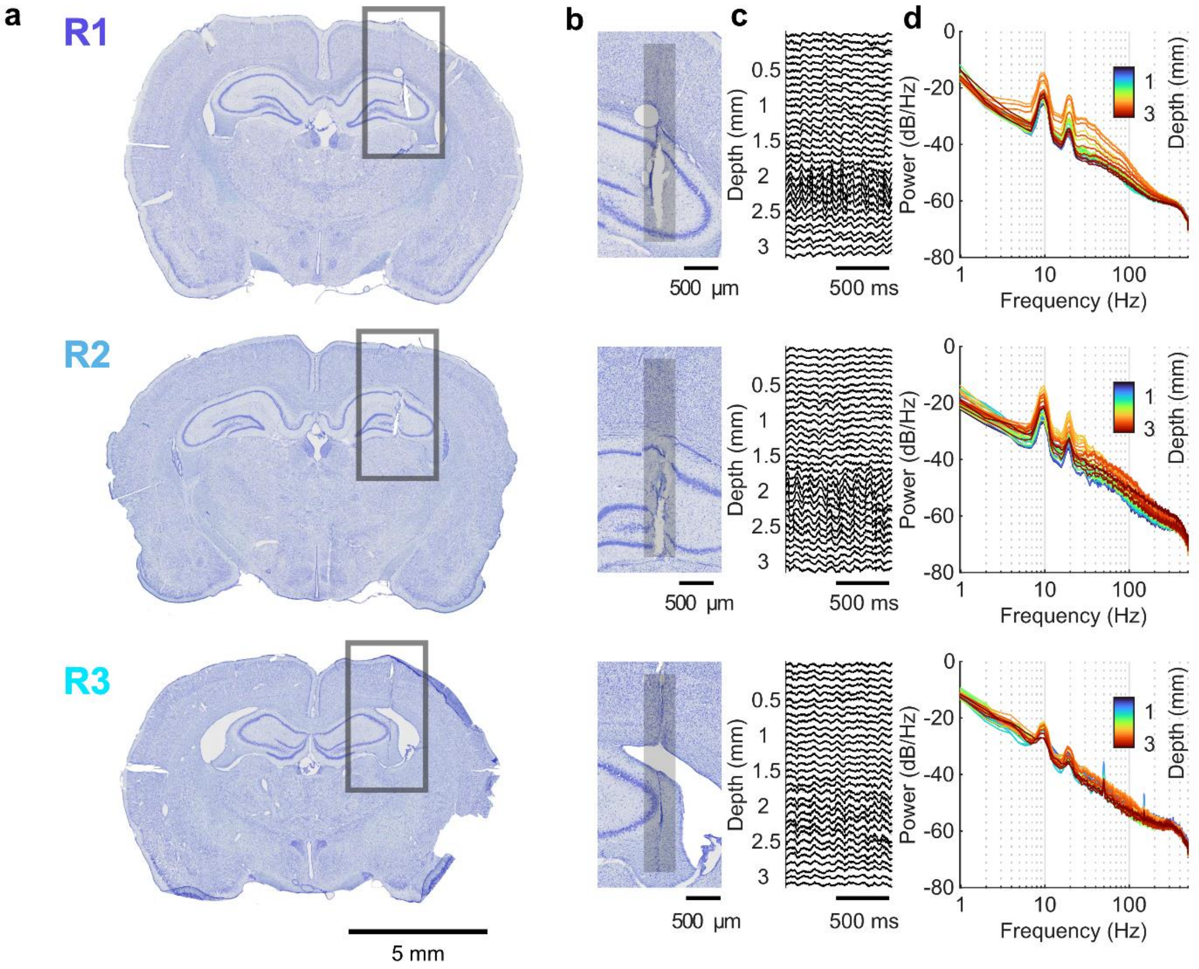


**Extended Data Figure 2: Rat electrode tracks and example LFP signals**

**a,** Nissl-stained coronal brain slices (40 µm thick) showing the track of the linear probes in rats R1, R2, and R3. **b,** Nissl-stained section of the hippocampus showing the estimated electrode track position (grey bar). **c,** LFP traces across all channels during locomotion (speed > 10 $cms^1$). **d,** Power spectral density of the LFP across all channels during locomotion for a single session.

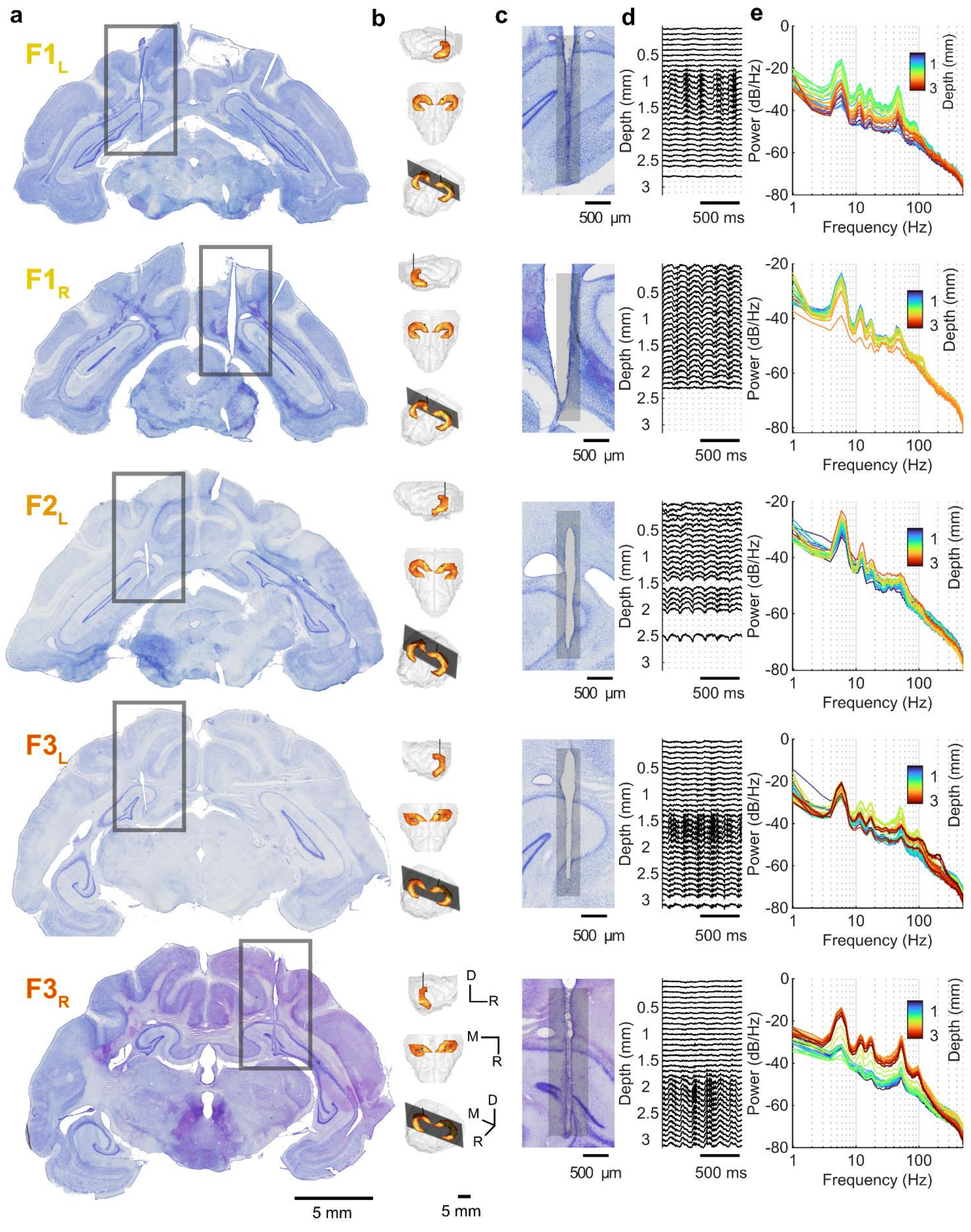


**Extended Data Figure 3: Ferret electrode tracks and example LFP signals**

**a,** Nissl-stained coronal brain slices (50 µm thick) showing the track of the linear probes in ferrets F1, F2, and F3. **b,** 3D reconstructions of each ferret brain. Black line indicates reconstructed electrode position. Grey planes indicate location of sections on the left. (D: dorsal, M: medial, R: rostral). **c,** Nissl-stained section of the hippocampus showing the estimated electrode track position (grey bar). **d,** LFP traces across all channels during locomotion (speed > 10 $cms^{-1}$). **e,** Power spectral density of the LFP across all channels during locomotion for a single session.

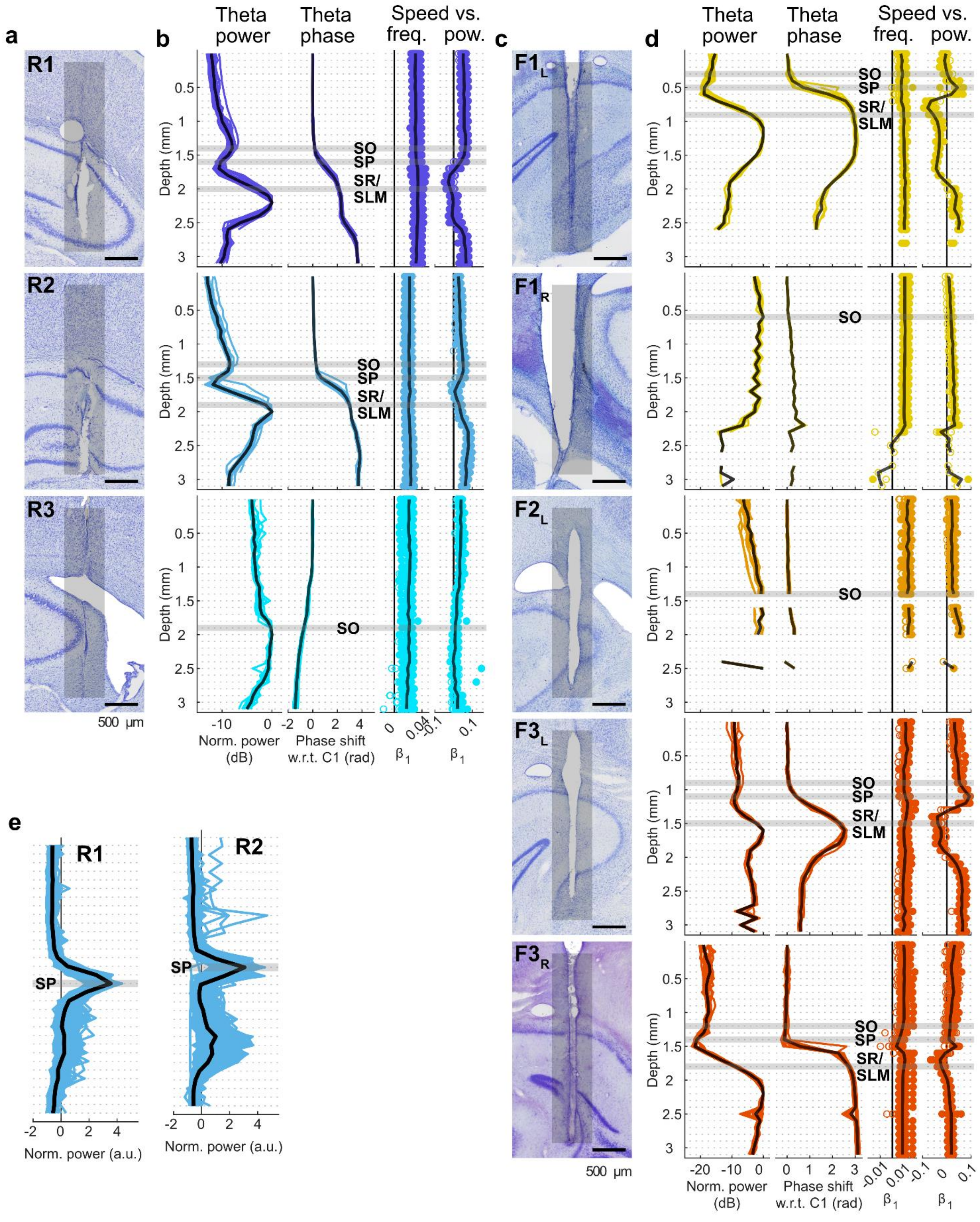


**Extended Data Figure 4: Theta depth profiles and hippocampal layer assignment**

**a,** Nissl-stained section of the hippocampus of rats R1, R2, and R3 showing the estimated electrode track positions (grey bars). **b,** Depth profile of theta characteristics across sessions. *Theta power:* power profile across the probe. Data shown as medians across probe, each line represents data from one session. *Theta phase:* phase shift for each channel relative to the top channel on the probe. Data shown as circular mean (CircStat Berens, 2009). *Speed vs freq.:* Regression slope of locomotion speed and theta frequency across the probe. *Speed vs pow.:* Regression slope of locomotion speed and theta power across the probe. Filled markers indicate significant regressions (Bonferroni corrected $p < 0.0016$). Estimated position of hippocampal layers based on theta depth profiles shown with grey shading (stratum oriens, SO; stratum pyramidale, SP, stratum radiatum/stratum lacunosum moleculare, SR/SLM). **c,** Nissl-stained section of the hippocampus of ferrets F1, F2, and F3 showing the estimated electrode track positions (grey bars). **d,** as in **b** but for ferrets F1, F2, and F3. **e,** Ripple power across the probes for rats R1 and R2, estimated position of the pyramidal cell layer based on the theta depth profiles shown with grey shading.

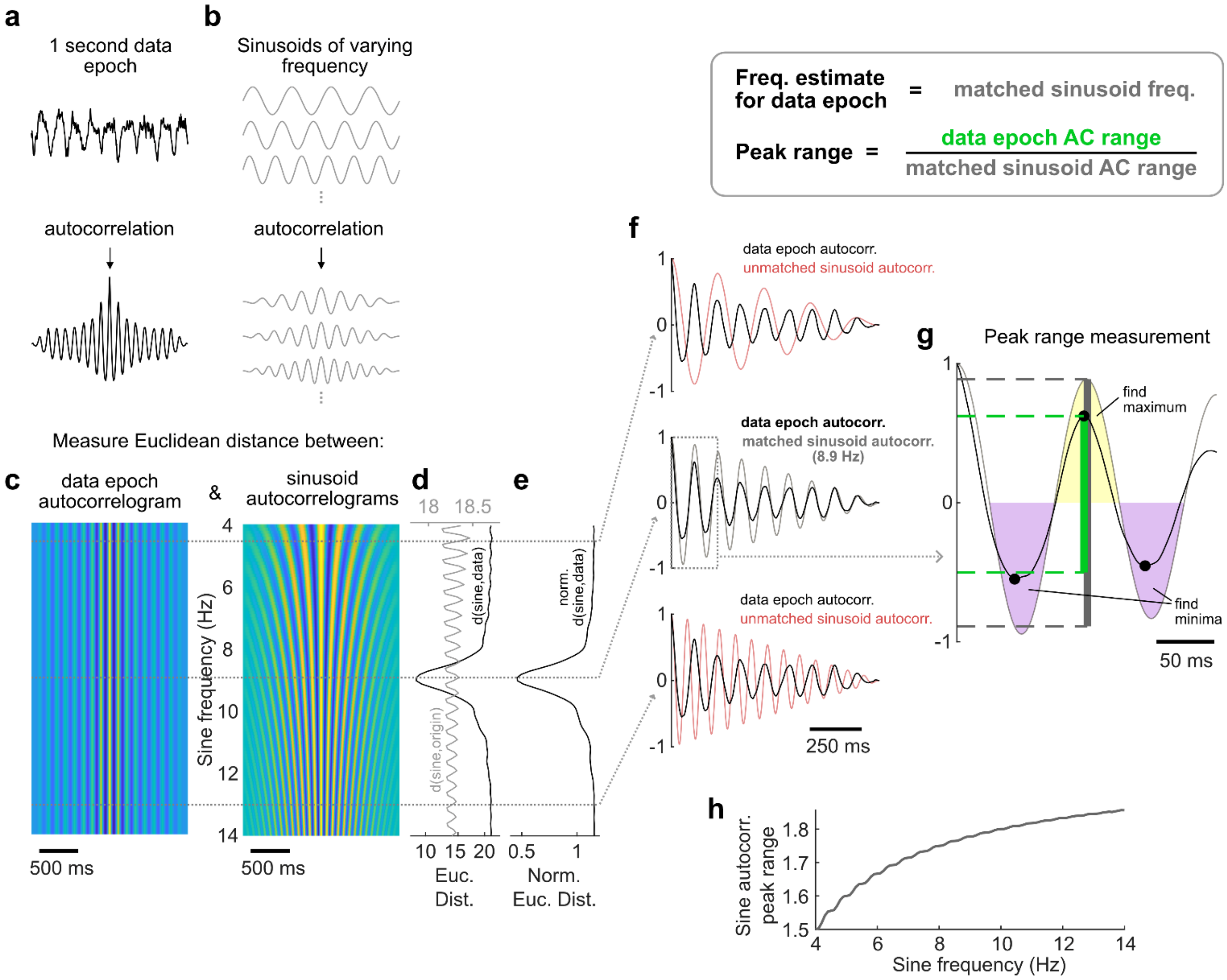


**Extended Data Figure 5: Autocorrelation quantification of oscillatory activity**

**a,** *Top:* Example 1 second LFP data epoch taken from rat R1 during locomotion. *Bottom:* Autocorrelogram of the data epoch above. **b,** *Top:* Example reference sinusoids. *Bottom:* Autocorrelograms of the example sinusoids above. **c,** The data epoch autocorrelogram visualised in a repeated manner (left) to illustrate the comparisons with the sinusoid autocorrelograms (right) across the full range of frequencies tested for the rat. **d,** Euclidean distance between the data epoch autocorrelogram and the sinusoid autocorrelograms (black). Grey line shows the Euclidean distance of the sinusoid autocorrelograms from the origin used for normalisation. **e,** Normalised Euclidean distance between the data epoch autocorrelogram and the sinusoid autocorrelograms. **f,** Examples of unmatched (top, bottom; red lines) and matched (middle; grey line) autocorrelograms with the data epoch autocorrelogram (black). **g,** Schematic of peak range measurement. The matched sinusoid (grey line) is used to define regions in which to search for the data autocorrelogram peak maxima (yellow shading) and minima (purple shading). The uncorrected peak range (green line) is calculated as the distance between the maxima to the mean of the minima, and is normalised by the peak range of the matched sinusoid autocorrelogram. **h,** The peak range of sinusoid autocorrelograms over frequency.

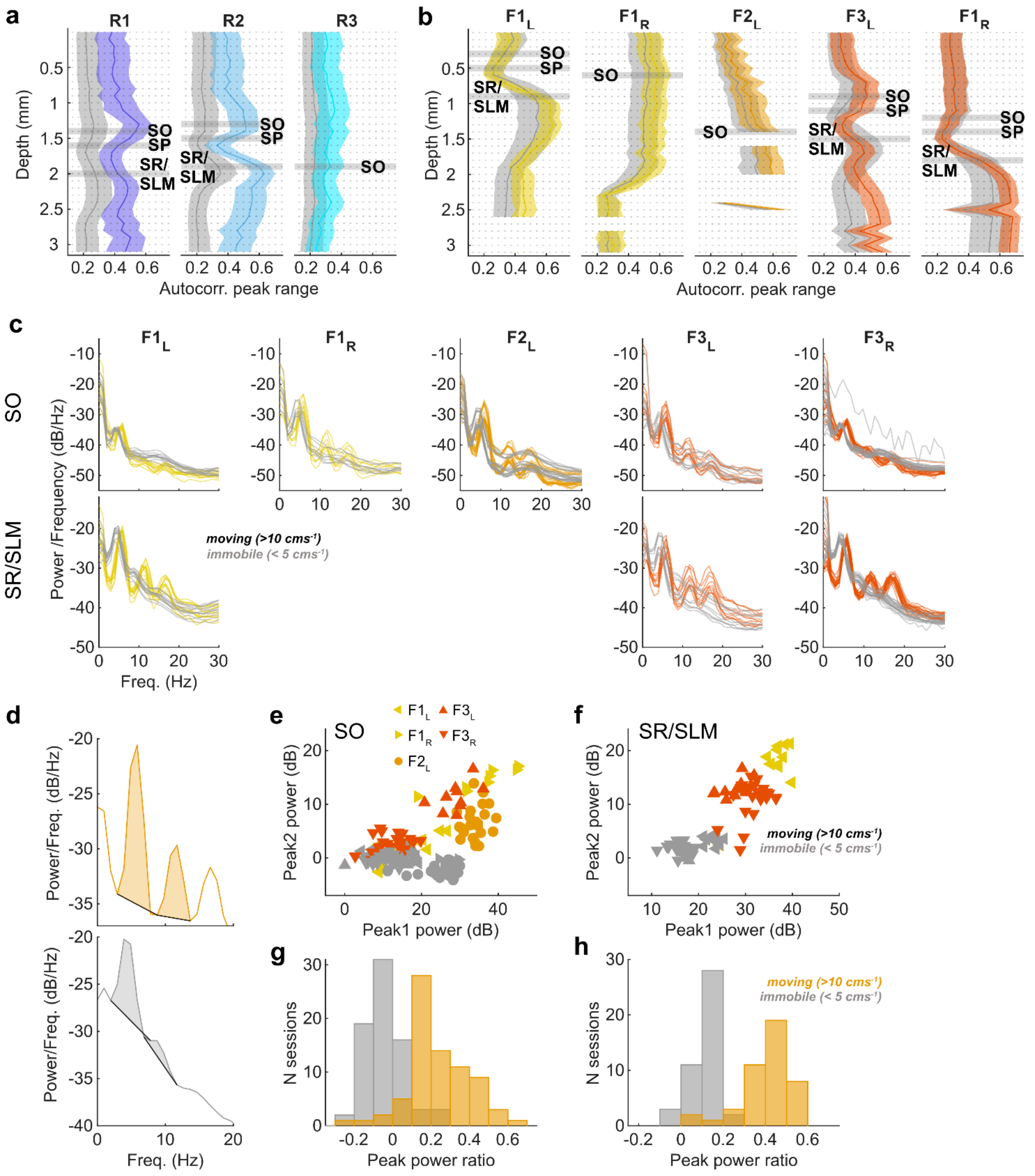


**Extended Data Figure 6: Depth profiles of peak range and PSD analysis of ferret LFP across locomotor contingency demonstrating that ferrets have robust low frequency oscillations during immobility with reduced saw-tooth profile**

**a,** Autocorrelogram peak range across all channels during locomotion (blue) and immobility (grey) for rats R1, R2, and R3. Estimated position of hippocampal layers based on theta depth profiles shown with horizontal grey shading (stratum oriens, SO; stratum pyramidale, SP, stratum radiatum/stratum lacunosum molecular, SR/SLM). **b,** As in **a** but for ferrets F1, F2 and F3; moving data is shown in orange. **c,** Power spectral density (PSD) analysis for moving (orange) and immobile (grey) data, recorded from the stratum oriens (OR; top row) and stratum radiatum/stratum lacunosum moleculare (SR/SLM; bottom row) of ferrets F1, F2, and F3. Each line represents data from one recording session. **d-h,** Quantification of the fundamental and first harmonic peaks in moving vs. immobile PSDs. **d,** Example PSD during locomotion (orange; top) and immobility (grey; bottom) illustrating the peak power estimation (shaded areas). **e-f,** Fundamental vs. first harmonic peak power for all ferret recording sites in the stratum oriens (SO; **e**) and stratum radiatum/stratum lacunosum moleculare (SR/SLM; **f**) during locomotion (orange) and immobility (grey). **g-h,** Power ratio of the fundamental and first harmonic peaks for all ferret recording sites in the stratum oriens (SO; **g**) and stratum radiatum/stratum lacunosum moleculare (SR/SLM; **h**) during locomotion (orange) and immobility (grey).

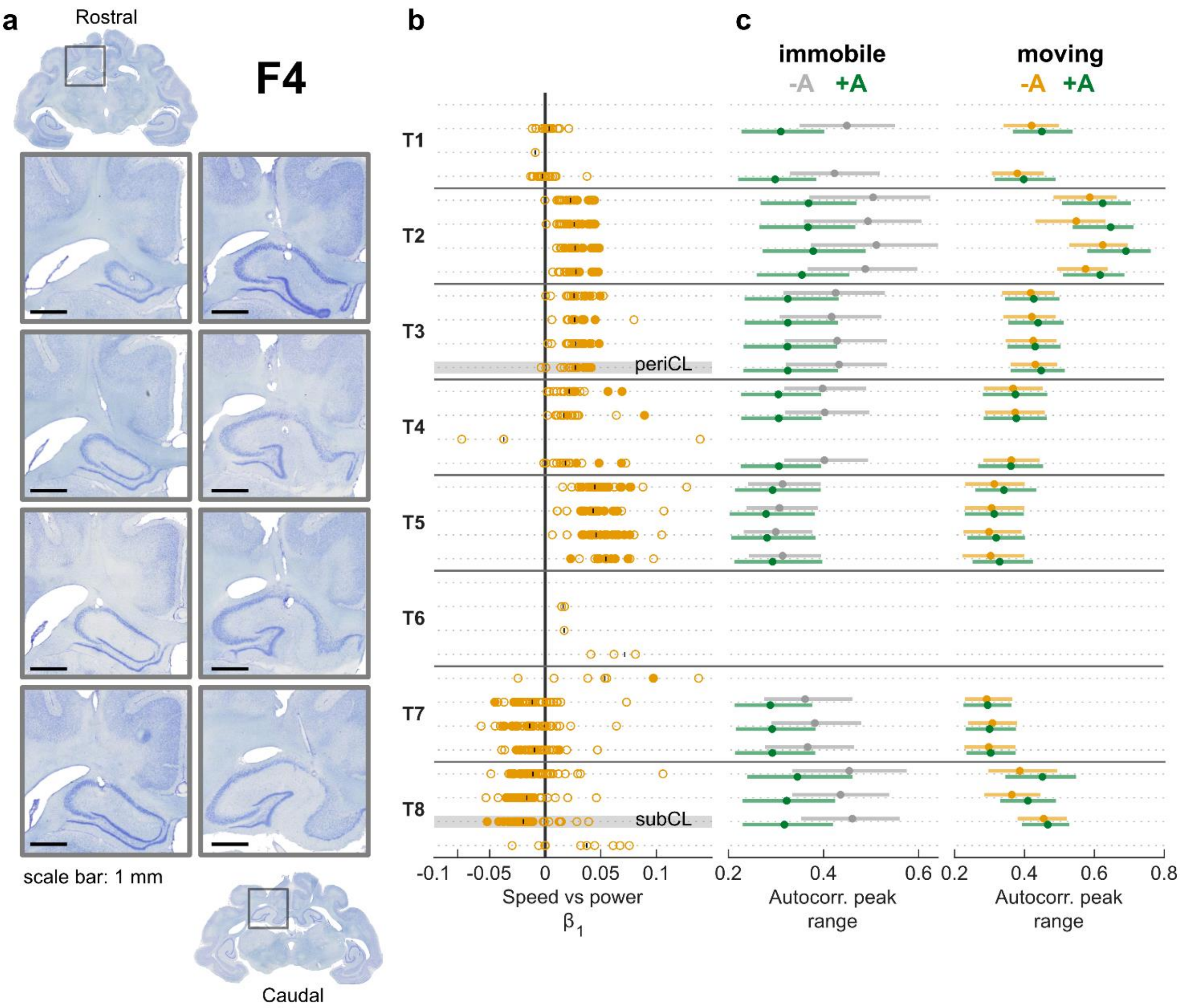


**Extended Data Figure 7: Histology, channel assignment, and effects of atropine administration across all channels in ferret F4**

**a,** Nissl-stained coronal sections (50 µm thick) showing electrode tracks and lesions in ferret F4. **b,** Regression slope of locomotion speed and theta power on every channel for all eight tetrodes (T1-8: T1-4 n = 15 sessions; T 4-8 n = 45 sessions). Each marker represents correlation calculated in one session. Filled markers indicate significant regressions (Bonferroni corrected $p < 0.0016$). Estimated position of hippocampal layers used in Figure 5 based on theta depth profiles shown with grey shading (periCL: defined by consistent positive speed-power correlations, subCL: defined by consistent negative speed-power correlations). **c,** Autocorrelation peak range during immobility (left) and locomotion (right) with and without the administration of atropine.

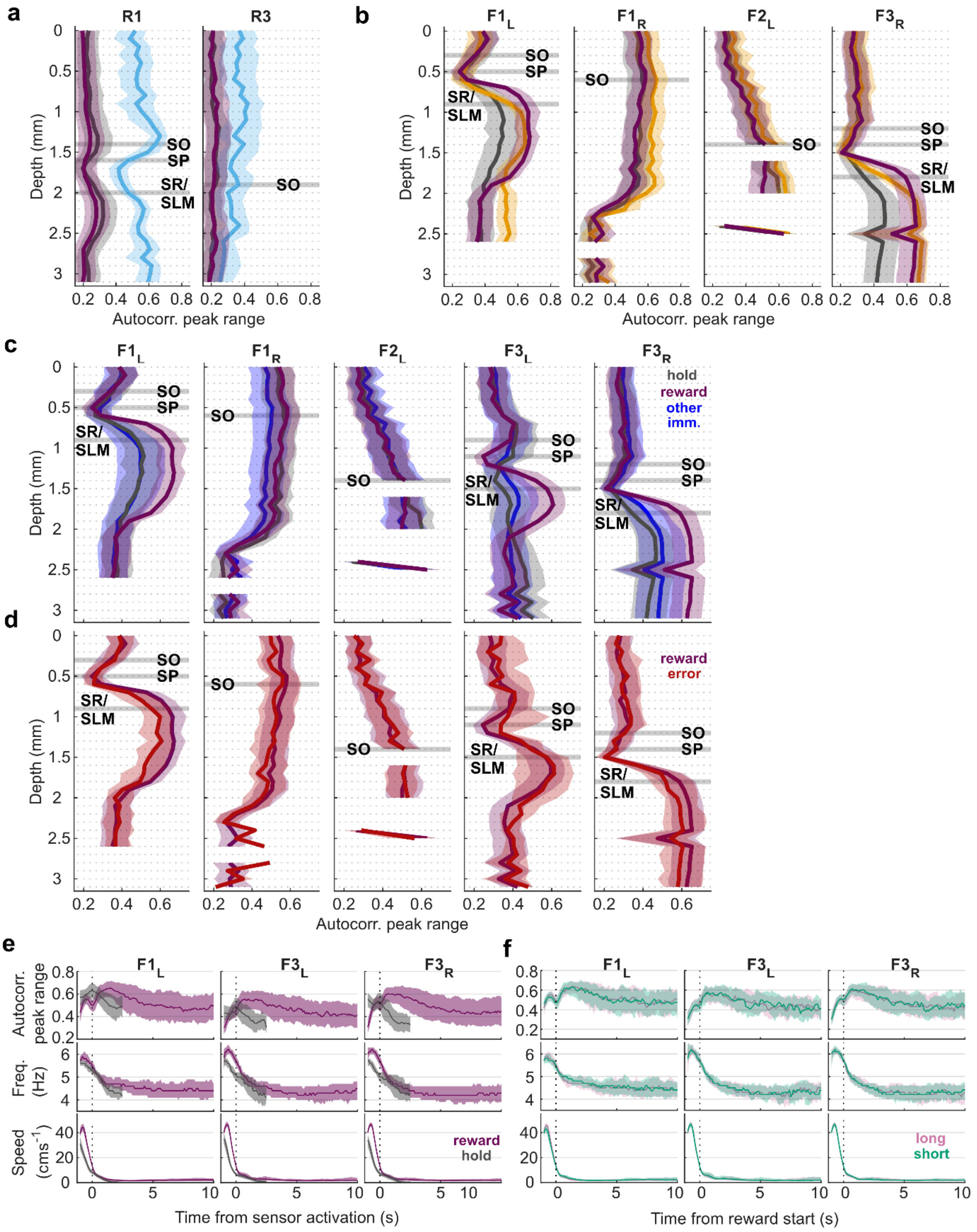


**Extended Data Figure 8: Oscillatory enhancement during Reward epochs in ferret stratum radiatum/stratum lacunosum moleculare are specific to peripheral spouts and not influenced by stimulus presentation**

**a,** Autocorrelation peak range for Hold (grey), Run (blue), and Reward (purple) epochs across all probe channels for rats R2 and R3; data are from all sessions and are presented as median (solid line) and interquartile range (shaded area). Channels in the stratum oriens (OR) and stratum radiatum/stratum lacunosum moleculare (SR/SLM) are highlighted with horizontal light grey shading. **b,** As in **a** but for ferrets F1, F2 and F3. Run data are presented in orange. **c,** Autocorrelation peak range during different periods of immobility (Hold, Reward, and Other; Other defined as immobility when the animal was not engaged in the task or at any spout locations) across all channels for ferrets F1, F2, and F3. Channels in the stratum oriens (OR) and stratum radiatum/stratum lacunosum moleculare (SR/SLM) are highlighted with horizontal light grey shading. **d,** Autocorrelation peak range for reward (i.e. correct) and error trials across all channels for ferrets F1, F2, and F3. Channels in the stratum oriens (OR) and stratum radiatum/stratum lacunosum moleculare (SR/SLM) are highlighted with horizontal light grey shading. **e,** Autocorrelation peak range (top), frequency (middle) and head speed (bottom) calculated using a sliding window (1 second) across reward at peripheral spouts (purple) and holding at the centre spout (grey) for channels in the ferret SR/SLM. **f,** Autocorrelation peak range (top), frequency (middle) and head speed (bottom) calculated using a sliding window (1 second) across reward at for trials with a long stimulus presentation (2 s; pink) and trials with a short (<1 s; teal) for channels in the ferret SR/SLM. This confirmed that there was no relation of reward enhancement to stimulus activation when behavioural responses overlapped with long-duration (2 s) stimuli.

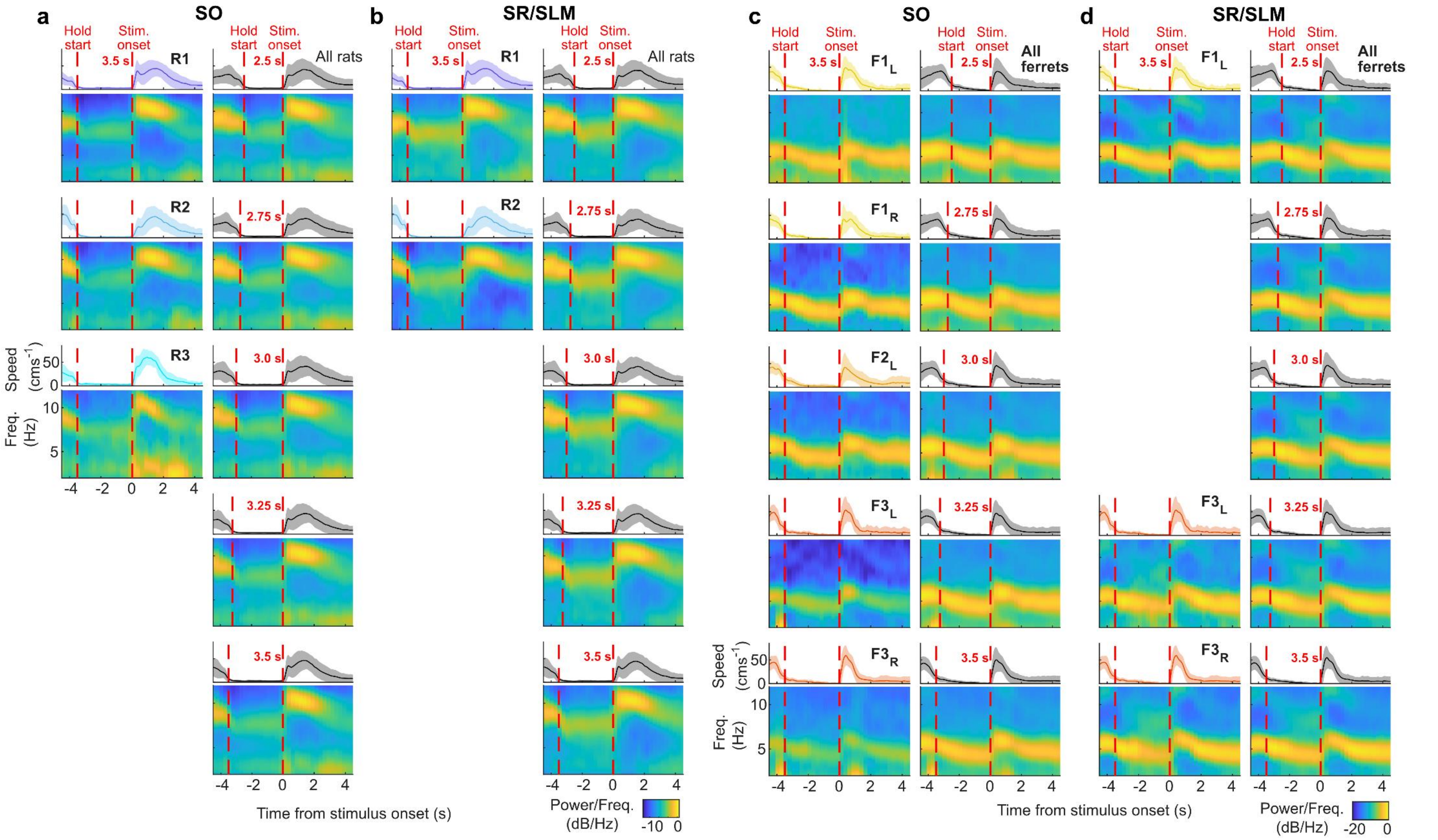


**Extended Data Figure 9: Mean trial spectrograms across individual implants and hold-times in the rat and ferret**
**a-b,** Mean speed traces and mean spectrograms of the LFP for all 3.5 s hold-time trials for each rat (left) and for all trials at each hold-time (left) for channels estimated to be in the stratum oriens (**a**; OR) and the stratum radiatum/stratum lacunosum moleculare (**b**; SR/SLM). Mean speed and standard deviation shown as line and shaded area. Red dashed lines indicate trial landmarks: the start of the hold where the animal initiates a trial and stimulus onset. **c-d,** As in **a-b** but showing ferret data.

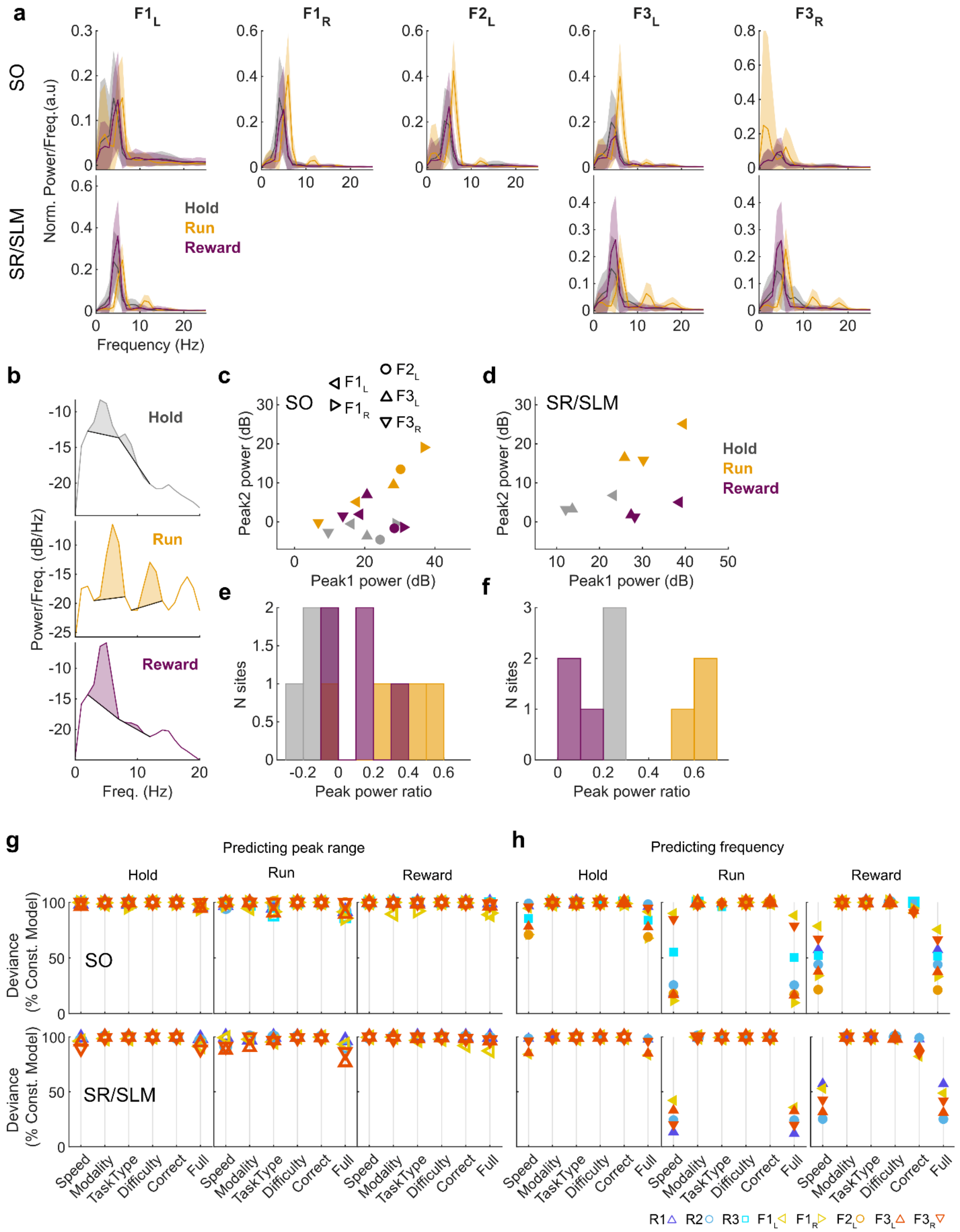


**Extended Data Figure 10: Theta during Hold and Reward epochs have similar wave shapes. No task parameters influence autocorrelation peak range measurements in any task epoch**

**a,** Power spectral density (PSD) of LFP from the three task epochs of interest (Hold: grey; Run: orange; Reward: purple) for each ferret in the stratum oriens (SO; top) and stratum radiatum/stratum lacunosum moleculare (SR/SLM; bottom). **b-f,** Quantification of the fundamental and first harmonic peaks in task epoch PSDs. **b,** Example PSD during Hold (grey; top), Run (orange; middle), and Reward (purple; bottom) illustrating the peak power estimation (shaded areas). **c-d,** Fundamental vs. first harmonic peak power for all ferret recording sites in the stratum oriens (SO; **c**) and stratum radiatum/stratum lacunosum moleculare (SR/SLM; **d**) during task epochs. **d-f,** Power ratio of the fundamental and first harmonic peaks for all ferret recording sites in the stratum oriens (SO; **g**) and stratum radiatum/stratum lacunosum moleculare (SR/SLM; **h**) during task epochs. **g,** Deviance of single-feature test models fitting autocorrelation peak range in each trial epoch expressed as a percentage of the deviance of the constant model. Full model shows performance of model including all predictors. Models fit separately for each individual rat and ferret. **h,** As in **g** but modelling frequency as the response variable.

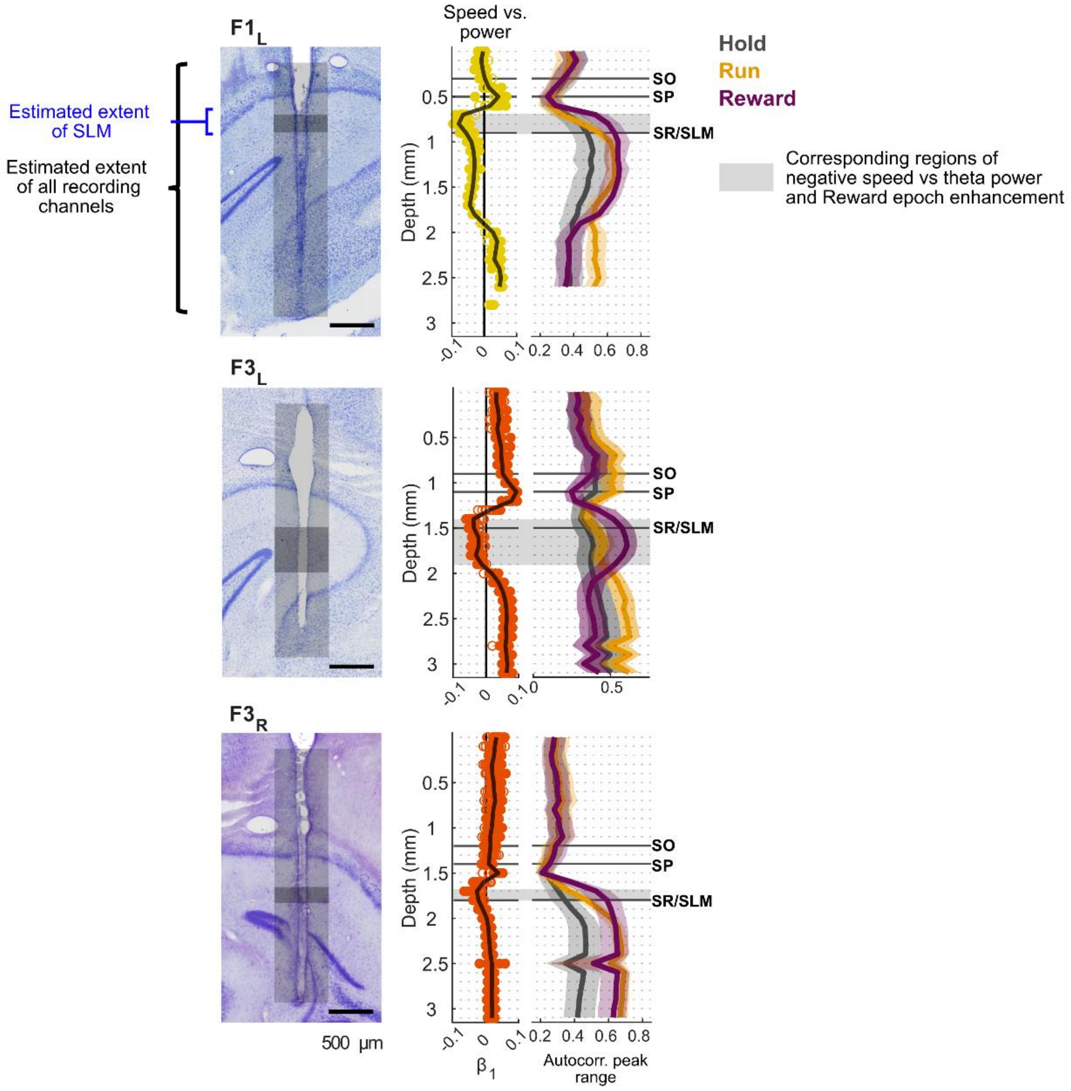


**Extended Data Figure 11: Reward epoch enhancement of immobility-related theta occurs on the same channels as negative speed-theta power correlations of locomotion-related theta, the extent of which appears to correspond with the distance that the probes travel through the stratum lacunosum moleculare**

*Left column:* Nissl-stained section of the hippocampus showing probe tracks for $F1_L$, $F3_L$, and $F3_R$. The estimated electrode track positions (light grey bars) and estimated extent of the stratum lacunosum moleculare (SLM; dark grey bars) are indicated. *Middle column:* Regression slope of locomotion speed and theta power across the probe. Filled markers indicate significant regressions (Bonferroni corrected $p < 0.0016$). *Right column:* Autocorrelation peak range for Hold (grey), Run (orange), and Reward (purple) epochs across all probe channels. Data are from all sessions and are presented as median (solid line) and interquartile range (shaded area). Estimated position of probe channels within hippocampal layers based on locomotion-related theta depth profiles shown with solid grey lines (stratum oriens, SO; stratum pyramidale, SP, stratum radiatum/stratum lacunosum moleculare, SR/SLM). Grey shaded regions in middle and left columns highlight the regions of negative speed-theta power correlations and Reward epoch enhancement of peak range.

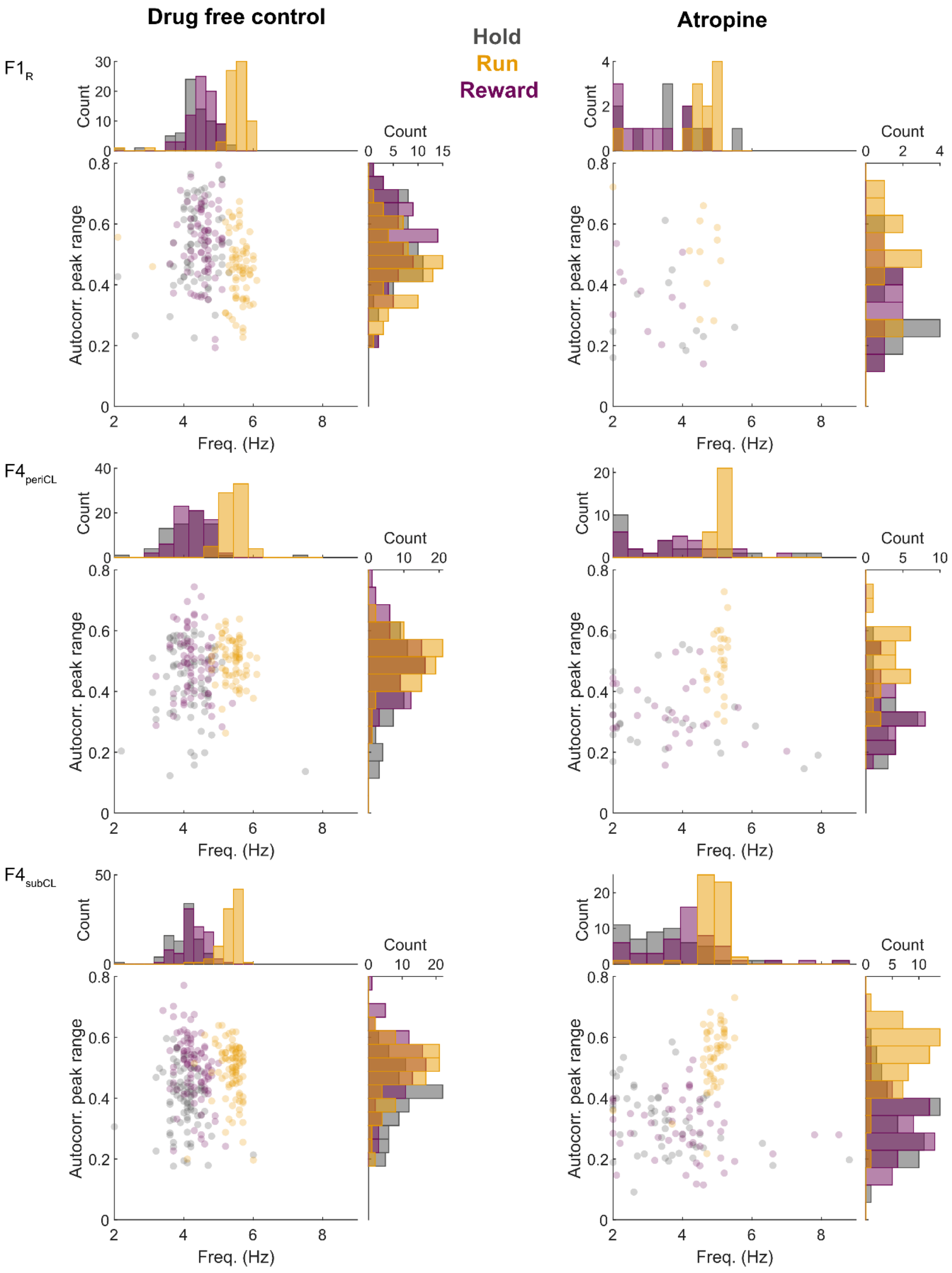


**Extended Data Figure 12: Theta in the Hold and Reward Epochs are similarly abolished by administration of atropine**
Scatter plot and marginal histograms showing relationship between LFP trial epoch frequency and normalised autocorrelation peak range for Hold (grey), Run (orange), and Reward (purple) epochs, for drug-free control sessions (left column) and during atropine administration (right column) for 3 channels across 2 ferrets (F1 and F4). Each marker represents data from a one second epoch from a single trial; data are from all sessions.
